# Glial immune-related pathways as mediators of closed head TBI effects on behavior in *Drosophila*

**DOI:** 10.1101/422535

**Authors:** Bart van Alphen, Samuel Stewart, Marta Iwanaszko, Fangke Xu, Eugenie Bang, Sydney Rozenfeld, Anujaianthi Ramakrishnan, Taichi Q. Itoh, Rosemary I. Braun, Ravi Allada

## Abstract

In traumatic brain injury (TBI) the initial injury phase is followed by a secondary phase that contributes to neurodegeneration. Yet the mechanisms leading to neuropathology in vivo remain to be elucidated. To address this question, we developed a *Drosophila* head-specific model for TBI, which we term Drosophila Closed Head Injury (dCHI), where well-controlled, non-penetrating strikes are directly delivered to the head of unanesthetized flies. This assay recapitulates many TBI phenotypes, including increased mortality, impaired motor control, fragmented sleep, and increased neuronal cell death. To discover novel mediators of TBI, we used glial targeted translating ribosome affinity purification in combination with RNA sequencing. We detected significant changes in the transcriptome at various times after TBI including in genes involved in innate immunity within 24 hours after TBI. To test the in vivo functional role of these changes, we examined TBI-dependent behavior and lethality in mutants of the master immune regulator NF-κB and found that while lethality effects were still evident, changes in sleep and motor function were substantially reduced. These studies validate a new head-specific model for TBI in *Drosophila* and identify glial immune pathways as candidate in vivo mediators of TBI effects.

Traumatic brain injury (TBI) is one of the leading causes of death and disability in the developed world [1-3]. Yet the underlying mechanisms that lead to long term physical, emotional, and cognitive impairment remain unclear.

Unlike in most forms of trauma, a large percentage of people killed by traumatic brain injuries do not die immediately but rather days or weeks after the insult [4]. TBI consists of a primary and a secondary phase. The primary brain injury is the result of an external mechanical force, resulting in damaged blood vessels, axonal shearing [5], cell death, disruption of the blood– brain barrier, edema, and the release of damage associated molecular patterns (DAMPs) and excitotoxic agents [6]. In response, local glia and infiltrating immune cells upregulate cytokines (tumor necrosis factor α) and interleukins (IL-6 and IL-1β) that drive post-traumatic neuroinflammation [7-10]. This secondary injury develops over a much longer time course, ranging from hours to months after the initial injury and is the result of a complex cascade of metabolic, cellular and molecular processes [11-13]. Neuroinflammation is beneficial when it is promoting clearance of debris and regeneration [14] but can become harmful, mediating neuronal death, progressive neurodegeneration, and neurodegenerative disorders [15-18]. The mechanisms underlying these opposing outcomes are largely unknown, but are thought to depend of the location and timing of the neuroinflammatory response [19, 20]. It remains to be determined what the relative roles of TBI-induced neuroinflammation and other TBI-induced changes are in mediating short and long-term impairments in brain function in vivo.

To study the mechanisms that mediate TBI pathology in vivo over time, we employ the fruit fly *Drosophila melanogaster*, a model organism well suited to understanding the in vivo genetics of brain injury. Despite considerable morphological differences between flies and mammals, the fly brain operates on similar principles through a highly conserved repertoire of neuronal signaling proteins, including a large number of neuronal cell adhesion receptors, synapse-organizing proteins, ion channels and neurotransmitter receptors, and synaptic vesicle-trafficking proteins [21]. This homology makes *Drosophila* a fruitful model to study neurodegenerative disorders [22], including ALS [23], Alzheimer’s disease [24], Huntington’s disease [25] and Parkinson’s disease [26].

Trauma-induced changes in glial gene expression are a highly conserved feature of both mammalian [27, 28] and *Drosophila* glia [29-32] (reviewed in [33]). In *Drosophila*, glia are able to perform immune-related functions [32, 34]. Ensheathing glia can act as phagocytes and contribute to the clearance of degenerating axons from the fly brain [29, 31, 35]. The *Drosophila* innate immune system is highly conserved with that of mammals and consists primarily of the Toll, Immunodeficiency (Imd) and Janus Kinase protein and the Signal Transducer and Activator of Transcription (JAK-STAT) pathways, which together combat fungal and bacterial infections [36, 37]. Dysregulation of cerebral innate immune signaling in *Drosophila* glial cells can lead to neuronal dysfunction and degeneration [38, 39], suggesting that changes in glia cells could underlie secondary injury mechanisms in our *Drosophila* model of TBI.

Existing *Drosophila* TBI models [40, 41] deliver impacts to the entire body, not just the head, and thus, one cannot definitively attribute ensuing phenotypes to TBI. To remove the confound of bodily injury, we have developed a novel, head-specific *Drosophila* model for TBI, *Drosophila* Closed Head Injury (dCHI). Here we show that by delivering precisely controlled, non-penetrating strikes to an unanesthetized fly’s head, we can induce cell death and increased mortality in a dose-dependent manner. In addition, TBI results in impaired motor control and decreased, fragmented sleep. Impaired motor control persists for many days after TBI while the sleep phenotype disappears after three days. These TBI-induced behavioral phenotypes do not occur in mutants lacking the master immune regulator NF-κB *Relish* (*Rel*), even though TBI-induced mortality is greatly induced in these mutants. In wild type flies, TBI results in changes in glial gene expression, where many immune related genes are upregulated 24 hours after injury. Together, these results establish a platform where powerful *Drosophila* genetics can be utilized to study the complex cascade of secondary injury mechanisms that occur after TBI in order to genetically disentangle its beneficial and detrimental effects.

## Methods

### Flies

Fly stocks were raised on standard cornmeal food under a 12h light/12h dark cycle at 25°C and ∼65% relative humidity. TBI inductions and climbing assays were carried out in the lab at room temperatures (∼ 21-23°C). For sleep and lifespan experiments, flies were kept on standard cornmeal food under a 12h light/12h dark cycle at 25°C and ∼65% relative humidity. All experiments were carried out in young adult *w*^*1118*^ males that are 3-7 days old. NF-κB Relish null mutants (Relish[E20]) were obtained from Bloomington (w^1118^; Rel^[E20]^ e^[s]^; #9457). Repo-Gal4 was obtained from Bloomington (w[1118]; P{w[+m*]=GAL4}repo/TM3, Sb[1] #7415). *UAS-GFP::RpL10A* was obtained from the Jackson lab [42]. All flies were collected under CO_2_ anesthesia at least 24 hours before TBI induction and placed on regular food.

### Aspirator and fly restraint assembly

Aspirators were constructed by wrapping a small square of cheesecloth around one end of aquarium tubing. A P1000 pipette tip was securely attached to covered end of the tubing, and the tip of the pipette tip was cut off to leave an aperture large enough for an individual fly to pass through without difficulty. The aspirator is used to transport individual flies via mouth pipetting. This allows flies to be transferred from their home vials to the experimental setup without using anesthesia. Fly restraints were created by cutting off the last 3-4 millimeters of P200 pipette tips to create an aperture large enough to let an individual fly’s head through without letting the entire body through. Multiple sizes of fly restraints were produced, to accommodate small variations in size among flies.

### Drosophila closed head TBI assay

Flies were removed from their home vials without the use of anesthetic, using an aspirator and gently transferred to a prepared P200 pipette (see above). By applying some air pressure on the aspirator, the fly is pushed into the P200 pipette in such a way that the fly gets stuck at the end, with only its head sticking out. The restrained fly is then placed in a micromanipulator allowing for movement in three dimensions, which was subsequently used to move the fly into the appropriate position, with the back of the fly’s head making contact with the pin of a pull-type solenoid (uxcell DC 12V) which delivers 8.34 Newtons of force. Flies were observed using a high-powered camera lens (Navitar Zoom 6000, Rochester, NY) to ensure that they were in the proper position. A variable-voltage power supply (Tenma Corporation, Tokyo, Japan) was set to 12V and used to power the solenoid, which then delivered a blow to the fly’s head (Fig. 1). Flies were hit 1 time, 5 times, and 10 times when observing effect of number of blows on response to TBI. Flies were hit 5 times for all other experiments. To minimize confounding effects of anesthesia, flies were collected under CO_2_ anesthesia at least 24 hours before each experiment. All experiments are carried out in awake, unanesthesized flies.

**Figure 1:**
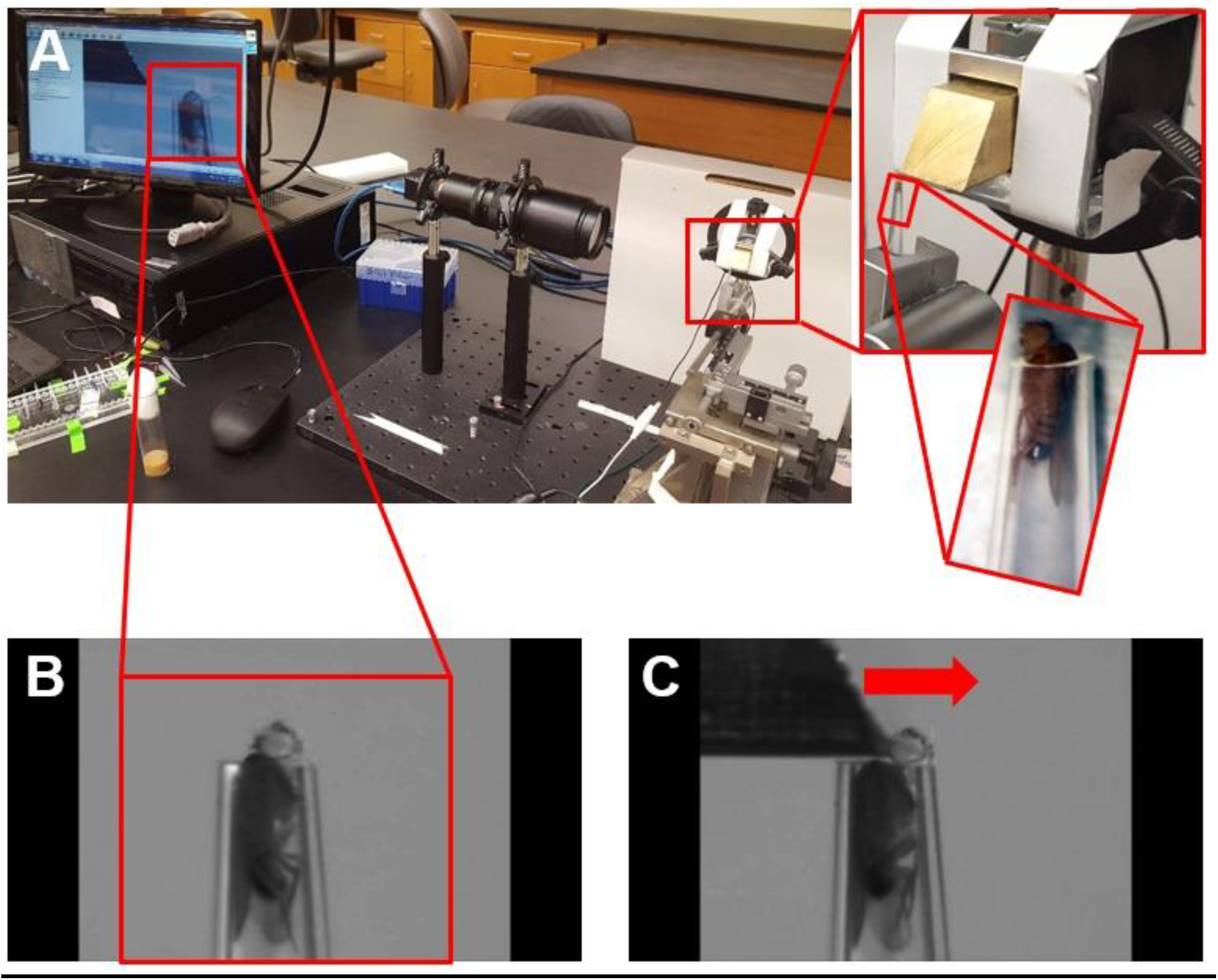
Single fly TBI setup. **A,B)** To induce head-specific TBI, individual flies are gently aspirated into a modified 200μl pipette tip that acts as a restraint. Immobilized flies are placed in front of a solenoid, using a set of micromanipulators with 5 degrees of freedom (x,y,z, pitch, roll) and a high magnification video system to ensure highly replicable positioning. TBI is induced by running a current through the magnetic coil of the solenoid, which retracts a brass trapezoid-shaped block. **C)** By releasing current, a spring drives the brass block forward, hitting the fly on the top of the head.

### Negative geotaxis response

A climbing assay is used to measure locomotor deficits after TBI in a manner similar to the RING assay [43]. Flies were individually stored in food vials and kept under the conditions discussed above. Vials were vertically divided into six 1-cm tall segments, labeled in order of ascending height (0 cm, 1 cm, etc.). Vials were tapped on a lab bench as a startle stimulus. Flies were then allowed to climb freely for 4 seconds, after which the highest point reached by the flies was observed and recorded. Three trials were observed for each individual fly; flies were allowed a period of at least 1 minute of recovery in between trials. Measurements from individual flies’ trials were then averaged to calculate a fly’s mean performance.

### Mortality assay

After TBI induction, flies are housed in plastic vials with standard corn meal medium and housed in a 12h light/12h dark cycle at 25°C and ∼65% relative humidity. Flies are gently transferred to fresh vials every three days. Deceased flies remaining in the old vial are counted.

### Sleep assay

3-7 days old flies were placed into individual 65-mm glass tubes in the Drosophila activity monitoring (DAM) system (Trikinetics, Waltham, MA), which were placed in incubators running a 12h light/12h dark cycle. All experiments were carried out at 25°C. Sleep data was collected by the DAM system in 1-minute bins and analyzed offline using custom made Matlab scripts (Matlab 2011a, Mathworks, Natick, MA). Briefly, sleep was defined as any period of inactivity of five minutes or more [44, 45]. For each fly, total amount of sleep per day, average bout length, number of sleep bouts, number of brief awakenings and average daily activity were derived from its activity trace (number of infrared beam crossings per minute).

### Statistics

All statistical analysis for behavioral experiments was performed using Matlab 2011a for PC. For TRAP-seq analysis, see below. Mortality assay: Survival curves were plotted using the Kaplan– Meier estimator as described [46]. The statistical significance was calculated using the log rank test. Plots and log rank tests were performed in Matlab, using scripts developed by [47].

### TUNEL assay

A TUNEL assay was performed in whole brain as per manufacturer’s protocol (In situ cell death detection kit, Fluorescein, Sigma Aldrich). The brains were carefully dissected out at different time points and fixed in 4% paraformaldehyde for 20 min followed by 3x 15 min wash in PBST (PBS with 0.5% Triton-X 100). The brains were incubated in TUNEL mixture (prepared as per manufacturer’s instruction) for 60 min at 37°C followed by 3 × 15 min wash in PBST. The brains were then mounted in Vectashield mounting medium.

### TRAP-seq

After receiving TBI, flies were collected at one of three time points, namely, 1 day post-injury, 3 days post-injury, and 7 days post-injury at ZT0 (lights-on in 12 h light: 12 h dark). Flies were collected in 15 ml conical tubes, and flash frozen in liquid nitrogen. Their heads were collected by vigorously shaking frozen flies and passing them through geological sieves. Approximately 100 heads were used for each experiment. Translating Ribosome Affinity Purification and Sequencing (TRAP-Seq) was performed as described [42, 48]. Sepharose beads were prepared by rinsing 25 μL of resin per reaction with 1 mL of extraction buffer. Protein A Plus UltraLink (PAS) resin was incubated with 1 mL of extraction buffer and 2.5 uG of HTZ 19C8 antibody and rotated for 2-3 hours at room temperature. Beads were then spun at 2500 g for 30 seconds at room temperature and rinsed another 3 times with extraction buffer. The conjugated beads were then incubated with 1 mL of blocking buffer for 15 minutes at 4°C. The beads were then spun again at 2500g for 30 seconds, and the supernatant was discarded. The beads were washed with 1 ml cold extraction buffer. This process was repeated another two times. Beads were incubated with 260 μL of head extract for 1 hour at 4°C, and then spun at 2500 g for 30 sec at 4°C. The beads were rinsed with 1 mL of cold wash buffer at 4°C. This process was repeated 3 times. After the final wash, 1 mL of Trizol was added. The beads were rotated at room temperature for 15 minutes. Chloroform was added, and the beads were subsequently shaken by hand for 30 seconds and incubated for 3 minutes at room temperature. The beads were then centrifuged at 15,000 rpm for 15 minutes at 4°C. The resulting upper aqueous phase was extracted and transferred to a new tube with 70% ethanol. RNA was extracted following the RNeasy Micro Kit protocol (Qiagen, Venlo, Netherlands). RNA purified from the GFP tagged RpL10 was then reverse transcribed to cDNA. The cDNA was used as template for T7 transcriptase to amplify the original RNA. Second round cDNA was synthesized based on amplified RNA. Detailed procedure for amplification can be in found in [49]. cDNA was sent to High Throughput Genome Analysis Core (HGAC) at the University of Chicago for library preparation and sequencing. Control and TBI samples were provided in three replicates.

### Quantification of data, differential expression and functional annotation analyses

Sequencing was done with Illumina HiSeq 2000. All samples are done with single end reads of 50 base pairs in length. At least ∼5,000,000 mappable reads were obtained and used for quantification for each sample. RNA-seq data were quantified at transcript level using Kallisto [50], with FlyBase_r6.14 as a reference transcriptome [51]. Quantified transcripts were summed up to the gene level using tximport library [52]. A minimal pre-filtering, keeping only rows with more than two reads, was applied to gene level data before differential expression (DE) analysis. Differential gene expression analysis was performed on TBI vs. control data with DESeq2 [53], using the likelihood ratio test to correct for batch effect among the biological replicates. Genes with the absolute log2 fold change higher than 0.6, and False Discovery Rate adjusted p-values ≤ 0.1 were identified as differentially expressed. Day 7 replicates were corrected for sequencing depth and possibly other distributional differences between lanes, using upper-quartile (UQ) normalization, available through RUVSeq library [54], before proceeding to DE analysis. One replicate was removed from further analysis, due to extremely low expression across the sample, which was not comparable to the levels observed in the other Day 7 replicates. Functional annotation of DE genes was performed using the DAVID database (release 6.8 [55, 56]) with a focus on gene ontology (GO) terms and Reactome pathways.

Supplemental Figures for post-TBI days 1, 3 and 7 (Fig. S2) show sample comparison of relative log expression in untreated and successfully corrected data (panels A, C, E and B, D, F respectively). Small deviations, arising from the technical differences, can be observed in D01 and D03, these were removed with upper-quartile between lane correction [57]. For D07 (Fig. S1E) we have observed that the replicates are lower quality and there is a significant deviation in values between replicates within the TBI group, with replicate R3 assumed to be corrupted (see Supplemental Table 2). For consistency we applied the same correction method to remove technical differences from post-TBI day 7, but as expected, replicate R3 did not improve. Taking this into consideration we decided to remove this replicate from further analysis.

## Results

### dCHI: A Novel Controlled Head Impact Model for TBI in Drosophila

To study TBI in flies, we developed a novel model where brain injury is inflicted in awake, individually restrained flies using a solenoid to deliver well-controlled, non-penetrating strikes to the fly head (Fig. 1). For TBI induction, individual flies are transferred from their home vial to a prepared P200 pipette tip, using an aspirator. Flies are gently blown upward until the head emerges from the tip of the pipette (Fig. 1B). The pipette is then placed in a micromanipulator platform with five degrees of freedom (pitch, roll as well as movement along the XYZ axes). The top of the fly head is pressed against the tip of the solenoid that consists of a metal pin running through a copper coil attached to a spring. By running a current through the coil, it acts as a magnet, drawing the pin back and arming the spring. When the current is halted, the spring causes the pin to shoot out, thus allowing us to deliver one or more blows to the fly’s head (Fig. 1C, Sup Movie 1). After TBI induction, flies are aspirated out of the pipette tip and returned to an empty vial containing regular fly food. Immediately after TBI induction, flies often seem dazed, being able to stand but only barely responding to tactile stimuli (Sup. Movie 2). However, mobility returns in a manner of minutes (Sup. Movie 3).

### dCHI increases mortality and impairs negative geotaxis in a dose-dependent manner within 24 h

To address the impact of dCHI, we first examined the acute pathological and behavioral effects within the first 24h post-dCHI. TBI phenotypes become more severe with consecutive strikes in mammals [58] and *Drosophila* [40, 41]. We subjected male flies to 1, 5 or 10 consecutive solenoid strikes, delivered at 1 strike per second. After TBI induction, treated and sham treated cohorts were individually housed in vials containing standard food. 24 hours after TBI exposure, surviving flies were counted in each of the four groups. We observed a dose dependent increase in 24-hour mortality (Fig. 2A). At 1 strike (TBIx1), there is no effect on 24-hour mortality (p=0.68). Mortality is increased in a dose dependent manner (control vs TBIx5, p = 0.03; control vs TBIx10, p = 0.004; ANOVA with Dunnett’s post hoc test, F(3,8) = 8.41; n = three replicates of 10 flies/group).

**Figure 2:**
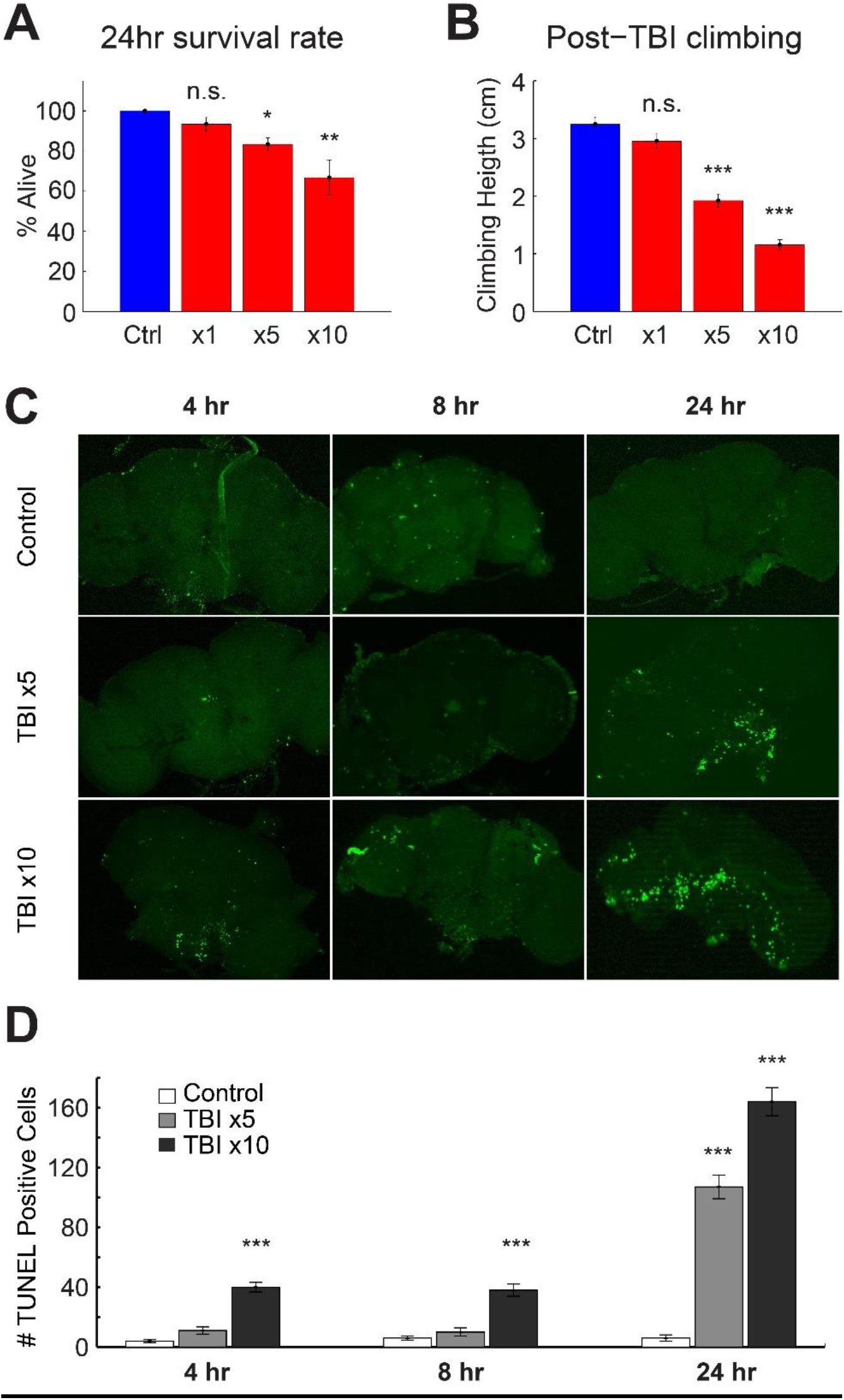
TBI causes cell death, mortality and impaired climbing in a dose-dependent manner. Male w^1118^ flies were exposed to either 1, 5 or 10 strikes to the head, delivered at 1 strike per second (n=32 per group). **A)** 24 hour survival rate decreased with increased number of strikes. **B)** in surviving flies, climbing behavior was quantified and compared to sham-treated controls 24 hours after TBI. Climbing behavior became more impaired with increased TBI severity (n.s = not significant, *** p < 0.001, one-way ANOVA with Dunnett post hoc test, n = 32/group). Cell death following TBI was quantified with a TUNEL assay **(C)** Representative images of TUNEL staining at different time points in control and post TBI flies **(D)** Histogram showing significantly increased TUNEL positive cells post TBI in a dose dependent manner (n = 10, 8, 8 for controls, n= 8, 8, 9 for TBIx5 and n = 7, 7, 8 for TBIx10 at 4hr, 8hr and 24 hr respectively). * p < 0.05, ** p < 0.01, *** p < 0.001 ANOVA with Dunnett posthoc test. Errorbars indicate SEM.

Loss of balance and poor motor coordination are symptoms associated with TBI [59-61]. Impairments in motor control, balance and sensorimotor integration are also a well-studied endophenotype in rodent models of TBI (as quantified by beam balance, beam walk and rotarod assays, reviewed in [62]). In *Drosophila*, impairments in sensorimotor integration are quantified by measuring the negative geotaxis response, a reflexive behavior where a fly moves away from gravity’s pull when agitated [63]. Impaired negative geotaxis has been observed in aging and in *Drosophila* models of neurodegeneration [64-66].

To assess sensorimotor function after TBI, we used a variation of the negative geotaxis assay [43], where the average height climbed in a defined time period is quantified, rather than a pass/fail number for absolute height as more subtle deficits can be observed using this approach. Typically, young adult wild-type flies reach an average climbing height of ∼4-5 cm in a 3-second time period [43]. In our assay, sham treated *w*^*1118*^ flies (3-7 days old males) reached an average height of ∼3.4 cm in four seconds (Fig. 2B, 0 days post-TBI). Climbing behavior, driven by negative geotaxis becomes impaired after TBI. After a single hit, there is no detectable difference in climbing, 24 hours after TBI induction (control vs TBIx1, p=0.1876, Fig. 2B). However, after five or ten consecutive hits, climbing behavior becomes impaired in a dose-dependent manner (Fig. 2B, control vs TBIx5, p = 2.67 × 10^−6^; control vs TBIx10, p = 2.67 × 10^−6^; ANOVA with Dunnett’s post hoc test, F(3,99) = 57.54; n = 30 flies/group)

### TBI increases apoptotic cell death in a dose and time-dependent manner

To test whether our TBI assay causes neuronal death, apoptosis was quantified using a TUNEL assay [67] after inducing TBI by striking flies either 5 or 10 times and comparing the number of TUNEL positive cells at three different timepoints (4, 8 and 24 hours) between TBI-treated flies and sham-treated controls. Controls showed, on average, two to four TUNEL positive cells which may be spontaneous apoptotic cells (Fig. 2C,D). Four hours after TBI induction we saw an increase in TUNEL positive cells in the TBIx10 condition (p = 2.56 × 10^−6^), but not in the TBIx5 condition (p = 0.1027; F(2,23) = 68.29) at this time point (Fig. 2D). Eight hours after TBI induction we also saw an increase in TUNEL positive cells in the TBIx10 condition (p = 2.93 × 10^−6^), but not in the TBIx5 condition (p = 0.5623; F(2,22) = 33.41) at this time point (Fig. 2D). 24 hours after TBI induction we saw an increase in TUNEL positive cells in both the TBIx5 (p = 2.57 × 10^−6^) and the TBIx10 condition (p = 2.53 × 10^−6^; F(2,19) = 111.23) at this time point (Fig. 2D). ANOVA with Dunnett’s post hoc test. Taken together, dCHI induces advanced mortality, motor deficits and cell death within the first 24 hours.

### dCHI reduces lifespan

Given the slowly evolving nature of TBI pathology, we next examined the chronic effects of dCHI over time. We first examined lifespan. Unlike other forms of trauma, death after TBI rarely occurs immediately. To test how our TBI assay affects overall lifespan, we delivered five consecutive strikes to the top of a fly’s head (Sup Movie 1). After this, flies were housed individually and survivors were counted every day. dCHI significantly reduces lifespan (log rank test on Kaplan-Meier survival curves, p < 0.001). 50% of the TBI group had died 14 days after TBI induction while 50% of the sham-treated controls had died 32 days after the start of the survival assay (Fig. 3A).

**Figure 3:**
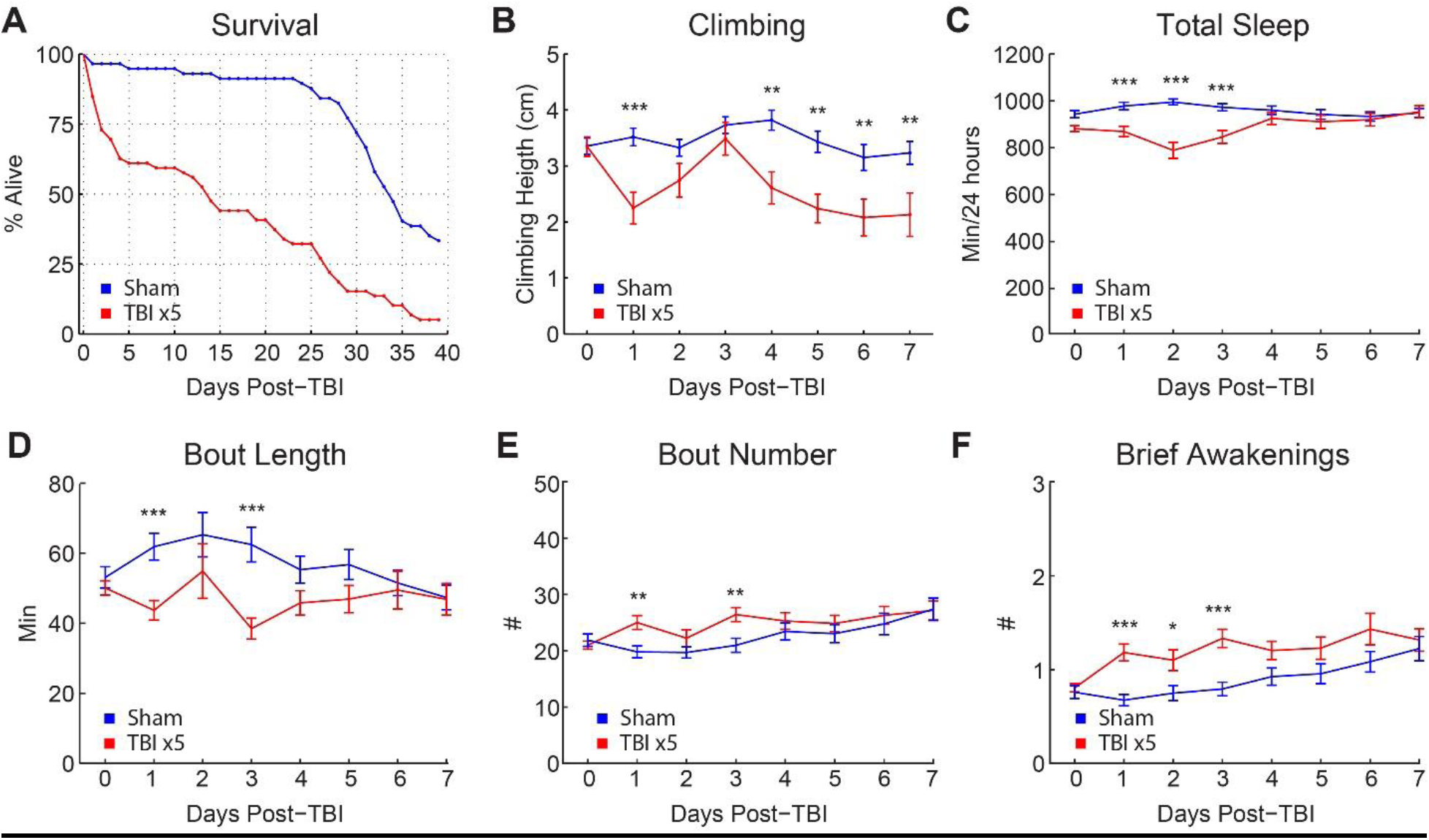
long term effects of TBI on mortality, climbing and sleep architecture. **A)** Kaplan-Meier estimates of survival functions in TBI treated flies and sham treated controls. TBI (5 strikes) was induced in male w1118 flies (n=59) and post-TBI survival was compared to survival in sham treated controls (n=57) using a log rank test. TBI results in a significant decrease in survival rate (p<0.001). 50% of the TBI group was deceased 14 days after TBI induction while 50% of the sham-treated controls had died 34 days after the start of the survival assay. **B)** Climbing behavior was tested in male w1118 flies., after which TBI was induced (n = 30). Climbing behavior was subsequently tested for seven days after TBI and compared to sham-treated controls (n = 30). Climbing impairments recover on post-TBI days 2 and 3, followed by a relapse on days 4-7 **C)** Sleep architecture was quantified in male flies up to ten days after TBI induction (n =103) and sham treated controls (n = 85). TBI induction resulted in **C)** decreased total sleep for up to three days post TBI, **D,E)** more fragmented sleep (decreased bout length, increased bout number) and **E)** increased brief awakenings, suggesting lighter sleep. *** p < 0.001, ** p < 0.01 by t-tests with Bonferroni correction). Errorbars indicate SEM.

### dCHI impairs motor control in a biphasic manner

One day after being subjected to five consecutive hits, injured flies display a decrease in climbing capacity, compared to sham-treated controls (Fig. 3B, p < 0.001, t-test with Bonferroni correction). Flies recover from days 2 to 3, suggesting that climbing deficits are not due to permanent injuries to the legs or to Johnston’s organ, the fly’s gravity sensor [68, 69]. Subsequently, flies undergo a relapse as climbing behavior is impaired again on days 4-7 (p < 0.01, t-tests with Bonferroni correction). The biphasic response to dCHI mirrors a similar biphasic motor response to TBI in a rodent model of TBI where rotarod performance was decreased at 2 and 30 days post TBI, but not at 7 days post TBI [70].

### dCHI reduces and fragments sleep

Sleep-wake disturbances after TBI are highly prevalent, occurring in 30-70% of TBI patients, and consisting of insomnia, hypersomnia, fragmented sleep and altered sleep architecture (reviewed in [71]). In rodent models of TBI, the most commonly reported sleep phenotypes are increased total sleep [72] [73-76] and increased sleep fragmentation [72, 73, 75, 77-79].

To test whether sleep is impaired in our TBI model, flies were individually loaded into Drosophila Activity Monitors immediately after TBI induction (5 strikes). The first three days post-TBI, sleep was reduced in TBI treated flies compared to sham treated controls (p < 0.001, t-test with Bonferroni correction, Fig. 3C). Also, sleep was fragmented, as seen by a decrease in the length of an average sleep bout (p < 0.001, t-test with Bonferroni correction, Fig. 3D) and an increase in the total number of sleep bouts (p < 0.01, t-test with Bonferroni correction, Fig. 3E). Brief awakenings, a measure of sleep depth [45] were increased (p < 0.001, t-test with Bonferroni correction, Fig. 3F). These sleep phenotypes were not evident after 4 days (Fig. 3C-F). Together, these results show that sleep in flies is decreased and fragmented in the first few days after TBI, but that it returns to baseline after four days.

### dCHI acutely activates the innate immune response

To test whether dCHI could alter glial gene expression, we used Translating Ribosome Affinity Purification and Sequencing (TRAP-seq). TRAP-seq allows for a cell type-specific analysis of the all mRNAs that ribosome associated and thus, are potentially being translated [48, 80].

We used the pan-glial driver *repo-GAL4* [81] to drive *UAS-GFP::RpL10A*, a GFP-tagged version of a ribosomal protein [48]. Head RNA was isolated at 1, 3 and 7 days after TBI induction and compared them to sham-treated controls. Ribosome associated mRNAs were isolated by immunoprecipiation of the RpL10A-GFP. For each time point, three replicates (n = ∼200 male fly heads/replicate) were collected. Gene expression levels were determined using Kallisto-derived estimated counts of glial TRAP-seq data collected at 1, 3, or 7 days after dCHI. Due to technical issues with one of day 7 TBI replicates, it was removed from further analysis (see Methods and Fig. S1).

To validate the method, we assessed enrichment of known glial genes in the anti-GFP immunoprecipitate (IP) relative to the input of whole head mRNAs (Fig. 4A). Expression levels for non-glia inputs were compared to glia-positive controls and show that glia-specific genes are enriched, including *astrocytic leucine-rich repeat molecule* (*alrm*)[29]) and *wunen-2* (*wun2*)[82], which are expressed highly in astrocytes, *gliotactin*, a transmembrane protein on peripheral glia [83]), *moody* (GPRCs that regulate blood brain barrier permeability [84]) and *reversed polarity* (*repo*) [81].

**Figure 4.**
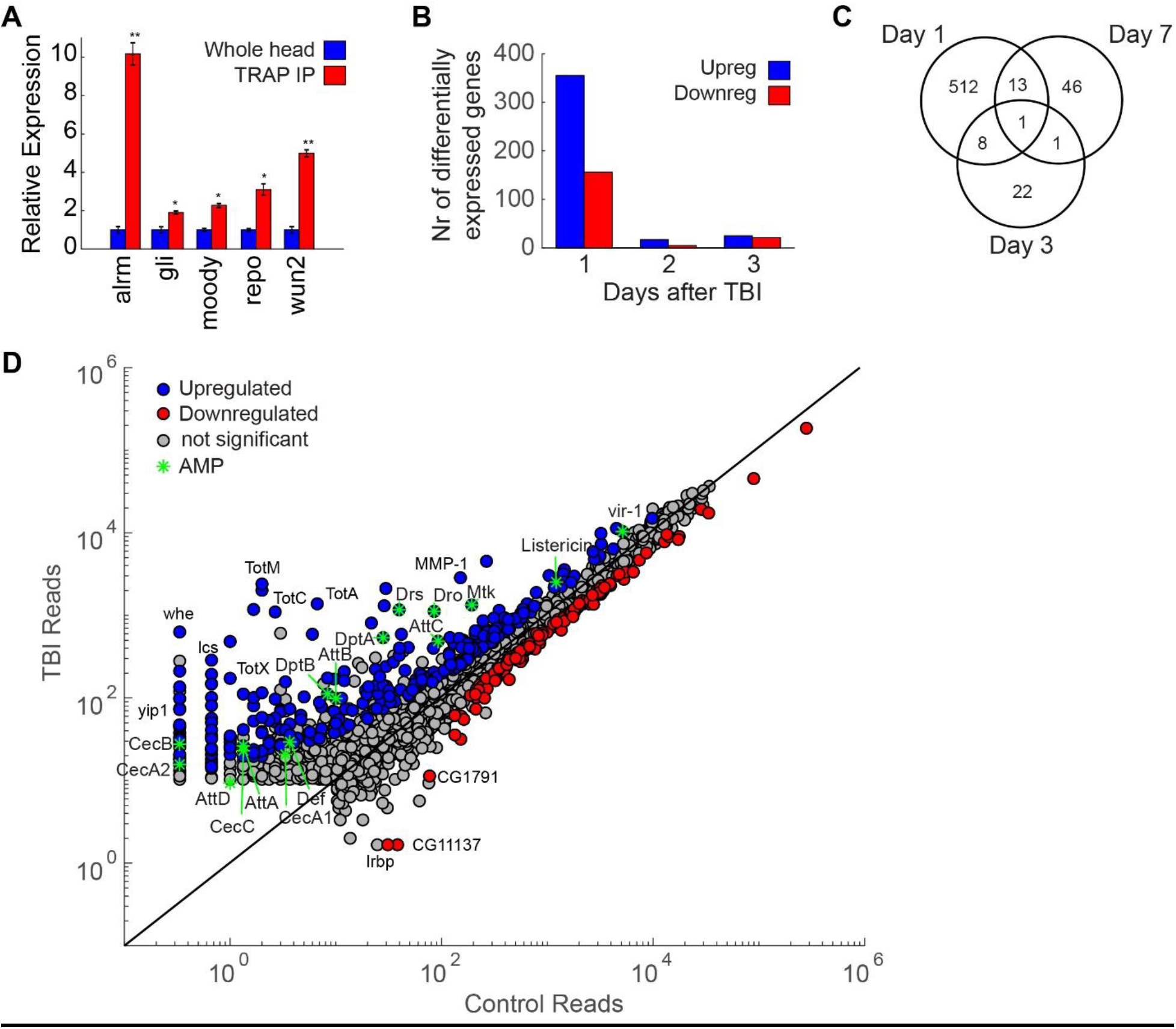
- TBI causes changes in gene expression. **A)** Expression levels for whole head inputs are compared to glia-specific TRAP immunoprecipitated (TRAP IP) and show that glia-specific genes are enriched, including astrocytic leucine-rich repeat molecule (alrm, astrocyte-specific), gliotactin (gli, expressed in peripheral glia), moody (blood brain barrier), reversed polarity (repo, pan-glial) and wunen-2 (wun2, astrocyte-specific). Data is normalized for whole head inputs (3 replicates for both whole head inputs and TRAP IP, ** p < 0.01, * p < 0.05, t-test). Errorbars show SEM. **B)** TBI induction results in a large number of differentially expressed genes in glia cells. 356 genes are upregulated and 156 are downregulated 1 day after TBI induction. Three days after TBI, gene expression is almost back to baseline, with only 17 genes that are upregulated and 5 genes that are downregulated. Seven days after TBI differential gene expression 24 genes are upregulated and 22 are downregulated. **C)** Venn diagram showing the overlap of differentially expressed genes (upregulated + downregulated) on 1, 3 and 7 days after TBI induction. **D)** Scatter plot for glial genes where average reads in the control condition are plotted against average reads 24 hours after TBI (blue dots, log_2_fold change ≥ 0.6, Benjamini adjusted p <0.1) or downregulated (red dots, log_2_fold change ≤ −0.6, Benjamini adjusted p <0.1) 24 hours after TBI induction. AMPs are indicated with green asterisks. Genes with average reads <10 in both control and TBI condition were excluded.

We initially looked at genes that are differentially expressed between dCHI and sham-treated controls. Log_2_ fold change shows how gene expression levels change after dCHI, compared to sham-treated controls. P-values are adjusted using the Benjamini-Hochberg procedure to correct for false discovery rates [85]. We first noted that the number of differentially expressed genes on day 1 post-dCHI vastly outnumber those evident on day 3 or day 7. In general, the number of upregulated genes is higher than the number of downregulated genes (Fig. 4B, Fig. S2). The differentially regulated genes are qualitatively different as well, with little overlap among differentially expressed genes on post-TBI day 1, 3 and 7 (Fig. 4C).

By plotting the average gene levels in the control group against those in TBI-treated flies (Fig. 4D), we see that TBI results in genes being more strongly upregulated than downregulated (more blue dots, further away from unity line). Strongly upregulated genes include antimicrobial peptides (AMPs) (Drosomycin (Drs), listericin, virus-Induced RNA-1 (vir-1), genes involved in proteolysis (alphaTry, yip7), and autophagy (MMP1). Also, quite a few members of the Turandot (Tot) family are upregulated after TBI (TotA, TotC, TotM, TotX). These genes are part of a broad stress response in *Drosophila* and are upregulated after exposure to mechanical stress, heat, UV, bacteria, oxidative stress and dehydration [86]. Turandot A (TotA) is strongly induced by bacterial challenge as well as exposure to mechanical pressure, dehydration and oxidative stress. [86]. We also find that several genes with poorly understood functions are also strongly upregulated after TBI (whatever (whe), la costa (lcs)).

The biological processes that are elevated 24 hours after TBI can be roughly grouped in three different categories: immune-related, proteolytic/protein folding and stress response processes. The majority of processes are part of the immune response (Table 1, red), including innate immune responses, humoral immune responses and different classes of antimicrobial peptides (AMPs). *Drosophila* AMPs can be grouped into three families based on their main biological targets, gram-positive bacteria (Defensin), gram-negative bacteria (Cecropins, Drosocin, Attacins, Diptericin), or fungi (Drosomycin, Metchnikowin) [87]. Most AMP genes were shown to be present in a TRAP-seq analysis of *Drosophila* astrocytes, a glial subset [88]. dCHI results in increased differential expression of many AMPs, including Attacins, Cecropins and Diptericins as well as Drosocin, Drosomycin and Metchnikowin (Fig. 5, blue bars). These AMPs are regulated by the Toll, Imd and JAK-STAT pathways [89]. Previous *Drosophila* TBI models showed an increase in three antimicrobial peptides (AMPs) after TBI induction, Attacin-C, Diptericin B and Metchnikowin [40, 41], but due to the non-specific nature of the impact, it is uncertain whether they are caused by TBI or by other types of injury. Enriched products of the antiviral and antibacterial JAK-STAT cascade are Listericin - an antibacterial protein and Virus-induced RNA 1, a marker of the induction of an antiviral response. In *Drosophila*, many proteases are involved in the immune response, including the activation of the Toll ligand Spätzle, which is under the control of a serine protease cascade [90]. These proteolytic cascades play a crucial role in innate immune reactions because they can be triggered more quickly than immune responses that require altered gene expression [91]. Expression of serine proteases that regulate Toll activation (Spirit, Grass, SME, Spheroid, Sphinx 1/2 [92]) were not significantly altered in our assay (data not shown). However, Späztle, a ligand for the Toll pathway that binds to Toll to activate the pathway [93], was upregulated. Activation of the immune response gradually dies down, as only three AMPs are upregulated three days after TBI. Seven days after TBI, all AMPs have returned to baseline levels (Fig. 5).

**Table 1:** Enriched biological processes 24 hours after TBI 24 hours after TBI, the Drosophila glial transcriptome shows a strong innate immune response as well as several physiological stress responses and proteolytic cascades.

| Term | Count | % | P-Value | Benjamini |
| --- | --- | --- | --- | --- |
| proteolysis | 55 | 10.8 | 6.40E-14 | 5.70E-11 |
| antibacterial humoral response | 13 | 2.6 | 4.70E-12 | 2.10E-09 |
| defense response to Gram-positive bacterium | 18 | 3.5 | 7.30E-12 | 2.20E-09 |
| response to bacterium | 16 | 3.1 | 4.00E-11 | 9.00E-09 |
| innate immune response | 22 | 4.3 | 1.30E-10 | 2.30E-08 |
| defense response | 11 | 2.2 | 2.30E-07 | 3.50E-05 |
| humoral immune response | 9 | 1.8 | 1.30E-06 | 1.70E-04 |
| protein folding | 16 | 3.1 | 8.60E-06 | 9.60E-04 |
| response to heat | 12 | 2.4 | 3.30E-05 | 3.20E-03 |
| defense response to Gram-negative bacterium | 15 | 3 | 3.80E-05 | 3.40E-03 |
| defense response to bacterium | 11 | 2.2 | 1.50E-04 | 1.20E-02 |
| protein refolding | 5 | 1 | 3.10E-04 | 2.30E-02 |
| rRNA pseudouridine synthesis | 4 | 0.8 | 4.60E-04 | 3.10E-02 |
| response to methotrexate | 5 | 1 | 7.00E-04 | 4.30E-02 |
| glutathione metabolic process | 8 | 1.6 | 1.50E-03 | 8.50E-02 |
| dorsal trunk growth, open tracheal system | 4 | 0.8 | 1.50E-03 | 8.20E-02 |
| cellular response to oxidative stress | 6 | 1.2 | 1.50E-03 | 7.80E-02 |
| chaperone-mediated protein folding | 5 | 1 | 1.80E-03 | 8.30E-02 |
| cytoskeleton organization | 7 | 1.4 | 1.80E-03 | 8.00E-02 |
| response to oxidative stress | 11 | 2.2 | 1.90E-03 | 8.30E-02 |
| cell adhesion | 11 | 2.2 | 2.30E-03 | 9.20E-02 |
| transmembrane transport | 22 | 4.3 | 2.30E-03 | 8.90E-02 |

- Immune response (115/286) - Protein folding/breakdown (85/286) - Physiological stress response (42/286) - Other

**Figure 5.**
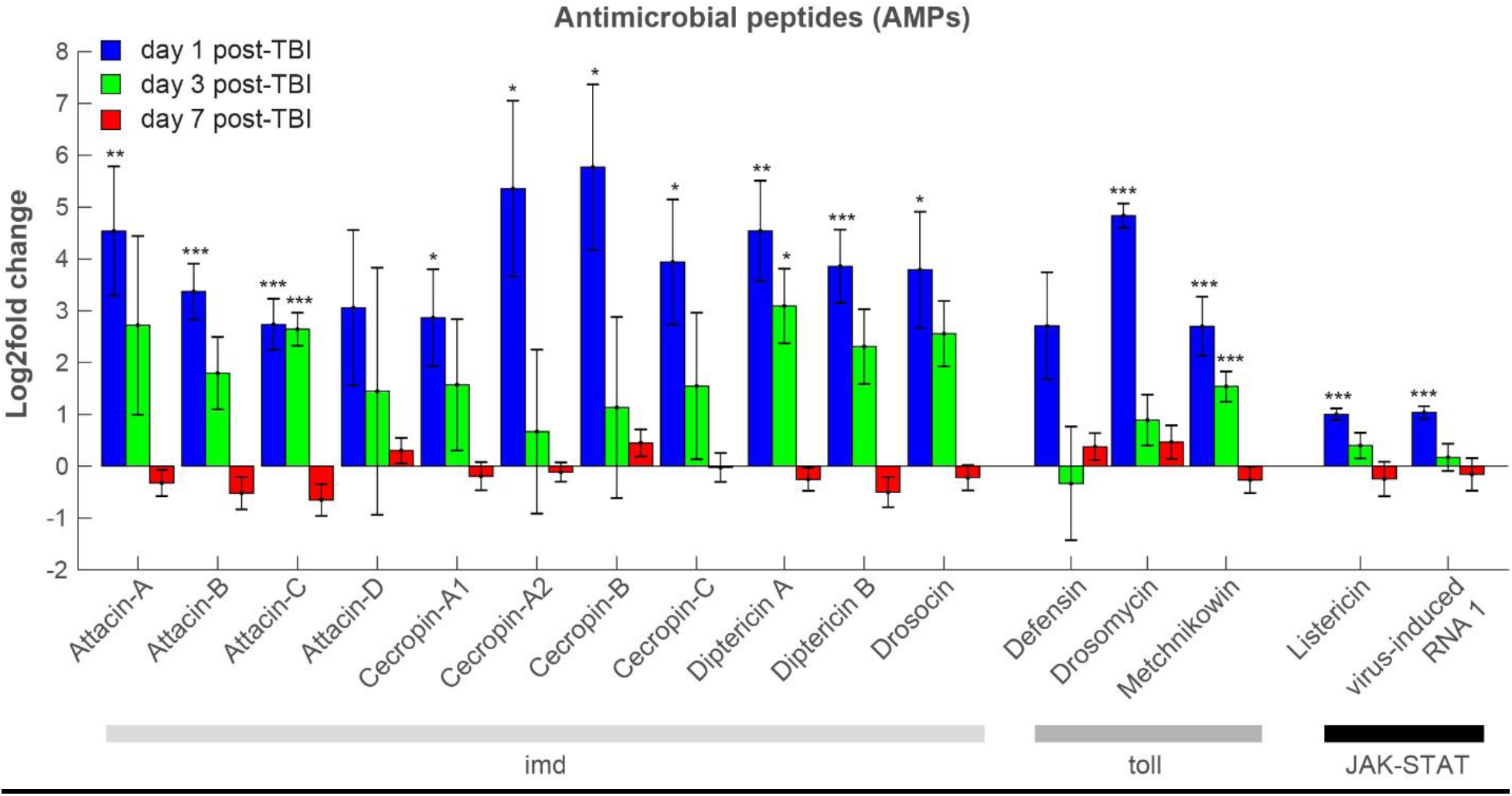
TBI activates a broad innate immune response. 24 hours after TBI, genes encoding most Drosophila antimicrobial peptides (AMPs) are enriched in glia cells. These include antibacterial, antifungal and antiviral peptides that are regulated by the Toll, Imd and JAK-STAT pathways. Three days after TBI (green), all but three AMPs (AttC, DptA, Mtk) have returned to control levels. Seven days after TBI (red) no AMPs are up- or down-regulated. Gene expression levels are considered to have changed significantly if the log2fold change ≥|0.6| and the Benjamini-adjusted p-value is <0.1. *** adj p < 0.001, ** adj p < 0.01, * adj p < 0.1. Errorsbars indicate SEM.

In addition to immune gene activation, transcriptomics also uncovered novel pathways that mediate TBI effects. A second category of enriched biological processes after TBI are proteases (Table 1). For example, we detected strong upregulation of matrix metalloproteinase-1 (MMP-1, 1.79 log_2_ fold change, adj p = 4.62E-23). MMPs are a family of endopeptidases that have diverse physiological and pathological functions, including degradation of extracellular matrix and regulation of cytokines/chemokines [94]. MMP-1 is induced in *Drosophila* ensheathing glia responding to axonal injury and is required for glial clearance of severed axons [35]. Detecting MMP-1 in our dCHI assay is further evidence that TBI damages axons and triggers glia-mediated neuroprotective responses.

The third main category of enriched genes 24 hours after TBI are stress response genes, genes that are upregulated in response to a variety of stressful stimuli, including heat, oxidative, metabolic and chemical stress [86, 95] (Table 1, green), including the Tot genes and heat shock proteins 22, 23, 26, 27, 68, 70Bc and lethal-2 as well as genes involved in oxidative stress. Heat shock protein 27 was previously implicated in response to bacteria and fungi as well [96].

In our model, TBI causes sleep to be reduced, fragmented and less deep (Fig.3C-F), a phenotype that persists for three days after TBI induction. Two prominent sleep regulating genes have altered expression levels 24 hours after TBI. Dopamine transporter (*DAT*, also known as *fumin*), a dopamine transporter that mediates uptake of dopamine from the synaptic cleft, is downregulated (−0.74 log_2_ fold change, p = 0.021). Loss of *DAT* increases extracellular dopamine and is associated with increased activity and decreased sleep [97-99]. *Pale* (*ple*), a tyrosine hydroxylase that drives synthesis of wake-promoting dopamine [68], is upregulated (0.63 log_2_ fold change, p = 0.018). *Pale* was previously shown to be activated in response to wounding in *Drosophila* embryos and larvae [100]. *Pale* and *DAT/fumin* levels are not changed on post-TBI days 3 and 7 (data not shown). Thus, we hypothesize down regulation of *DAT/fumin* in combination with upregulation of *pale* may underlie TBI changes to sleep due to increased dopamine levels.

Three days after TBI induction, there are few significant differences in gene expression between TBI treated flies and sham treated controls (Fig. 4B, Fig. S3A). Whereas there are 512 genes with altered expression 24 hours post-TBI (almost 400 of those are upregulated), after three days there are only 22 genes with altered expression levels (Fig. 4B). Interestingly, this low level of glial activation at day 3 post-TBI coincides with a climbing behavior returning back to control levels (Fig. 4B). At three days post TBI, several AMPs remain strongly upregulated (AttC, Mtk, Dpt; Fig. 5, Fig. S3A).

Seven days after TBI induction, there is more variability in gene expression (Fig. S1E,F). There seems to no activation of the immune response, as all AMPs have returned to baseline levels (Fig. 5, red bars). There is only one gene that is persistently downregulated on days 1, 3 and 7 (Fig. 4C). CG40470’s function is unknown, although roles in proteolysis and peptide catabolic processes have been inferred [101].

### Post-TBI behavioral phenotypes are NF-κB -dependent

The NF-kB family of transcription factors plays a central role in the regulation of inflammatory gene expression [102], cell survival and neuronal plasticity [103]. NF-kB is activated in neurons and glial cells after injury and has been linked to both neurodegenerative and neuroprotective activities [103, 104]. NF-kB mediates activation of glial cells [105] and inflammation [106]. TBI causes an increase in Nuclear Factor kappa B (NF-κB) in rodents [107-109], where it has a neuroprotective effect in a closed-head model of TBI [70]. In *Drosophila*, NF-κB is a crucial component of both the Toll and the Imd pathways, where different isoforms are required for the expression of different antimicrobial peptides [110]. Changes to sleep architecture after injury/infection in *Drosophila* require NF-κB *Relish* [111]. In *Drosophila*, over-expression of NF-κB or AMPs in glia cells causes neurodegeneration [38, 39, 112, 113].

**Fig 6.**
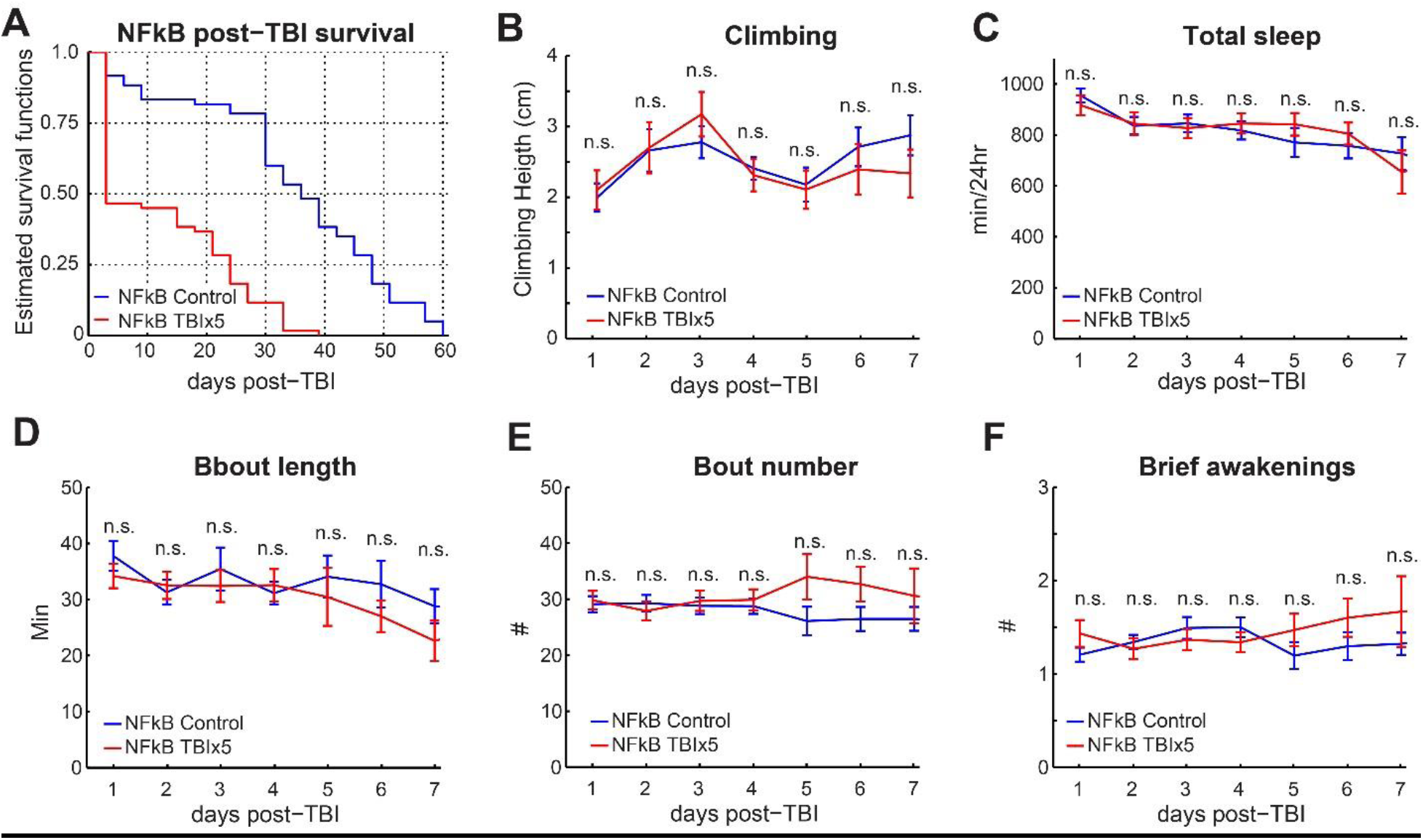
Post-TBI behavioral phenotypes are NF-kB dependent. Kaplan-Meier estimates of survival functions in TBI treated NF-κB null mutants and sham treated controls (n=60/group) show that TBI significantly reduces lifespan (log rank test, p <0.001). **B)** The effect of TBI on climbing behavior was tested in male NF-κB null mutants and sham-treated controls (n = 32/group for seven consecutive days after TBI induction. There was no significant difference between both groups (n.s., t-tests with Bonferroni correction. **C)** TBI does not impair sleep in NFκB null mutants, compared to sham-treated controls. TBI does not fragment sleep as average bout length **(D)** and bout number **(E)** are not changed. **F)** Brief awakenings, a measure of sleep depth, are unchanged after TBI. (n = 52 controls, 57 in TBI group, n.s.; t-tests with Bonferroni correction). Errorbars indicate SEM.

To test the role of NF-κB in TBI-induced mortality and behavioral impairments, we induced TBI in the NF-κB *Relish* null mutant (w1118; Rel[E20], Bloomington #9457) and measured its effects on post-TBI survival, climbing behavior and sleep. Five consecutive strikes to the head resulted in strongly increased mortality (log rank test on Kaplan-Meier survival curves, p < 0.001), where over 50% of the NF-κB mutants had died three days after TBI induction (Fig. 6A). In the background strain (*w*^*1118*^), 50% mortality for five strikes occurs at ∼14 days post-TBI (Fig.3A). In sham-treated NF-κB controls, 50% mortality occurs after 35 days (Fig. 6A), which is very similar to mortality in the sham-treated *w*^*1118*^ background strain (Fig. 3A).

To test whether TBI has a much stronger effect on NF-κB mutants, we tested climbing behavior daily for seven days after TBI induction. However, there was no difference between TBI treated flies and untreated controls on any day (Fig. 6B). This is different from wild-type flies which show considerable impairment to climbing 24 hours post-TBI, which then reverts back to normal and is followed by a relapse on days 5-7 (Fig. 3B). Both TBI treated and sham treated controls show a gradual decrease in total sleep over ten days post-treatment. However, there are no differences in total sleep (Fig. 6C) or in any other metrics of sleep architecture (brief awakenings, bout length, bout number, wake activity; Fig. 6D-F). These results suggest that the NF-κB-dependent immune response facilitates survival after TBI but that impairments in sleep and climbing behavior are consequences of an immune-dependent injury mechanism.

### Discussion

Here we have developed a simple and reproducible *Drosophila* model for closed head TBI where we deliver precisely controlled strikes to the head of individually restrained, unanesthetized flies. This TBI paradigm is validated by recapitulating many of the phenotypes observed in mammalian TBI models, including increased mortality, increased neuronal cell death, impaired motor control, decreased/fragmented sleep and hundreds of TBI-induced changes to the transcriptome, including the activation of many antimicrobial peptides, indicating a strong activation of the immune response. Behavioral responses to TBI (e.g. sleep and geotaxis) are abolished in mutants of the transcription factor NF-κB *Relish*, which plays a central role in regulating stress-associated and inflammatory gene expression in both mammals [103, 114] and flies [115]. Nonetheless, *Relish* null mutants show increased mortality after TBI. These results set the stage to leverage *Drosophila* genetic tools to investigate the role of the immune response as well as novel pathways in TBI pathology.

Our single fly paradigm is a more valid *Drosophila* model for TBI that circumvents the lack of specificity of currently available models [39, 41]. Both previous assays induce TBI by subjecting the whole fly to trauma, which makes it hard to distinguish whether observed phenotypes are a due to traumatic brain injury or a consequence of internal injuries. A recently published method [116] uses a pneumatic device to strike an anesthetized fly’s head. This method is an improvement of earlier assays and results in increased mortality in a stimulus strength dependent manner. However, it only shows a modest reduction in locomotor activity, without demonstrating any other TBI-related phenotypes such as neuronal cell death or immune activation. The dependence on CO_2_ anesthesia further impairs the usefulness of this assay, as prolonged behavioral impairments in *Drosophila* occur even after brief exposure to CO_2_ anesthesia [117]. Additionally, anesthetics that are administered either during or shortly after TBI induction can offer neuroprotective effects and alter cognitive, motor, and histological outcomes in mammalian models of TBI [118-120] as well as affecting mortality in a whole-body injury model in flies [121]. Our *Drosophila* model allows us to study how TBI affects behavior and gene expression without the confounding effects of anesthesia, making it a more valid model for TBI that occurs under natural conditions.

In this study, we also elucidate in an unbiased manner, the genomic response to TBI. Glial cells play an important role in immune responses in both mammals and *Drosophila*, (reviewed in [33]). Until now, profiling TBI-induced changes in gene expression have either been limited to a small number of pre-selected genes in both mammals [122, 123] and *Drosophila* [40, 41] or focused on whole brain tissue rather than individual cell types [124, 125]. Using TRAP in combination with RNA-seq, we detect an acute, broad-spectrum immune response, where antimicrobial peptides (AMPs) and stress response genes are upregulated 24 hours after TBI. These include antibacterial, antifungal and antiviral peptides as well as peptides from the Turandot family which are secreted as part of a stress response induced by bacteria, UV, heat and mechanical stress [86]. Three days after TBI, only Attacin-C, Diptericin A and Metchnikowin are upregulated. Seven days after TBI, AMPs or stress response genes are not detectably upregulated. These findings match reports in mammalian TBI models, where inflammatory gene expression spikes shortly after TBI but mostly dies down during subsequent days [122, 126].

Besides validating our *Drosophila* model with the detection of a strongly upregulated immune response, we detected several novel genes among the total of 512 different glial genes that were either up- or down-regulated after TBI. Immune and stress response only make up 157 out of 512 differentially expressed glial genes. Genes involved in proteolysis and protein folding are a prominent portion (85/512) of these differentially expressed genes, yet their role in TBI is poorly understood. These results demonstrate that there are other candidate pathways that may modulate recovery, and *Drosophila* can be used as a first line screen to test their in vivo function.

We have successfully applied in vivo genetics to identify in vivo pathways important for TBI. Here we demonstrate that loss of master immune regulator NF-κB results in increased mortality after TBI, yet it protects against TBI-induced impairments in sleep and motor control. These findings align with previous reports showing links between sleep and the immune response in flies [127] where NF-κB is required to alter sleep architecture after exposure to septic or aseptic injuries [128]. One possibility is that sleep impairments can be a side-effect of melanization, an invertebrate defense mechanism that requires dopamine as melanin precursor [129]. If dopamine is upregulated to create more melanin, decreased sleep would be a side effect. Consistent with this hypothesis, we observe changes in *fumin* and *pale* which could results in increased dopamine levels. Recently, it was shown that repressing neuronal NF-κB in a mouse model of TBI increases post-TBI mortality, as in our studies, without reducing behavioral impairments [70], suggesting that non-neuronal NF-κB could underlie behavioral impairments after TBI. The demonstration of an in vivo role for TBI-regulated genes will be important for defining their function.

In summary, our *Drosophila* Closed Head Injury assay (dCHI) recapitulates many of the physiological symptoms observed in mammals, demonstrating that fruit flies are a valid model to study physiological responses to TBI. We demonstrate both a potent induction of immune pathways and a requirement for an immune master regulator in mediating TBI effects on behavior. Our model now provides a platform to perform unbiased genetic screens to study how gene expression changes after TBI in unanesthetized, awake animals result in the long-term sequelae of TBI. These studies raise the possibility of rapidly identifying key genes and pathways that are neuroprotective for TBI, thereby providing a high-throughput approach that could facilitate the discovery of novel genes and therapeutics that offer better outcomes after TBI.

## Supplemental Figures

**Fig S1.**
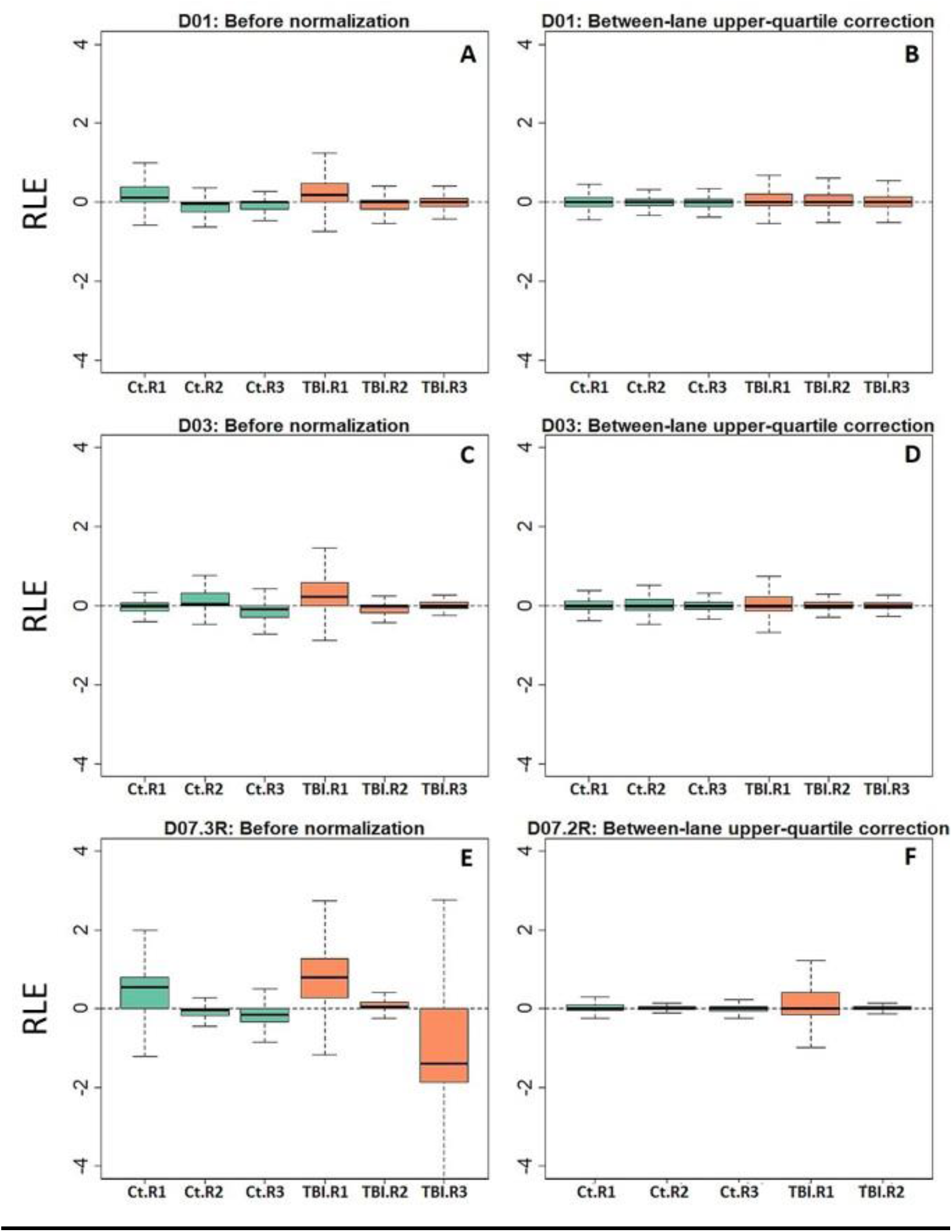
Relative log expression and normalized data for post-TBI days 1, 3 and 7. Relative log expression (RLE) plot of raw and normalized glial expression data. Control (green) and TBI (orange) biological replicates for day 1, 3 and 7 post-TBI. Correction was performed using the upper-quartile between lane method. Due to the high variability in TBI replicate 3 on day 7, this replicate was discarded

**Fig S2:**
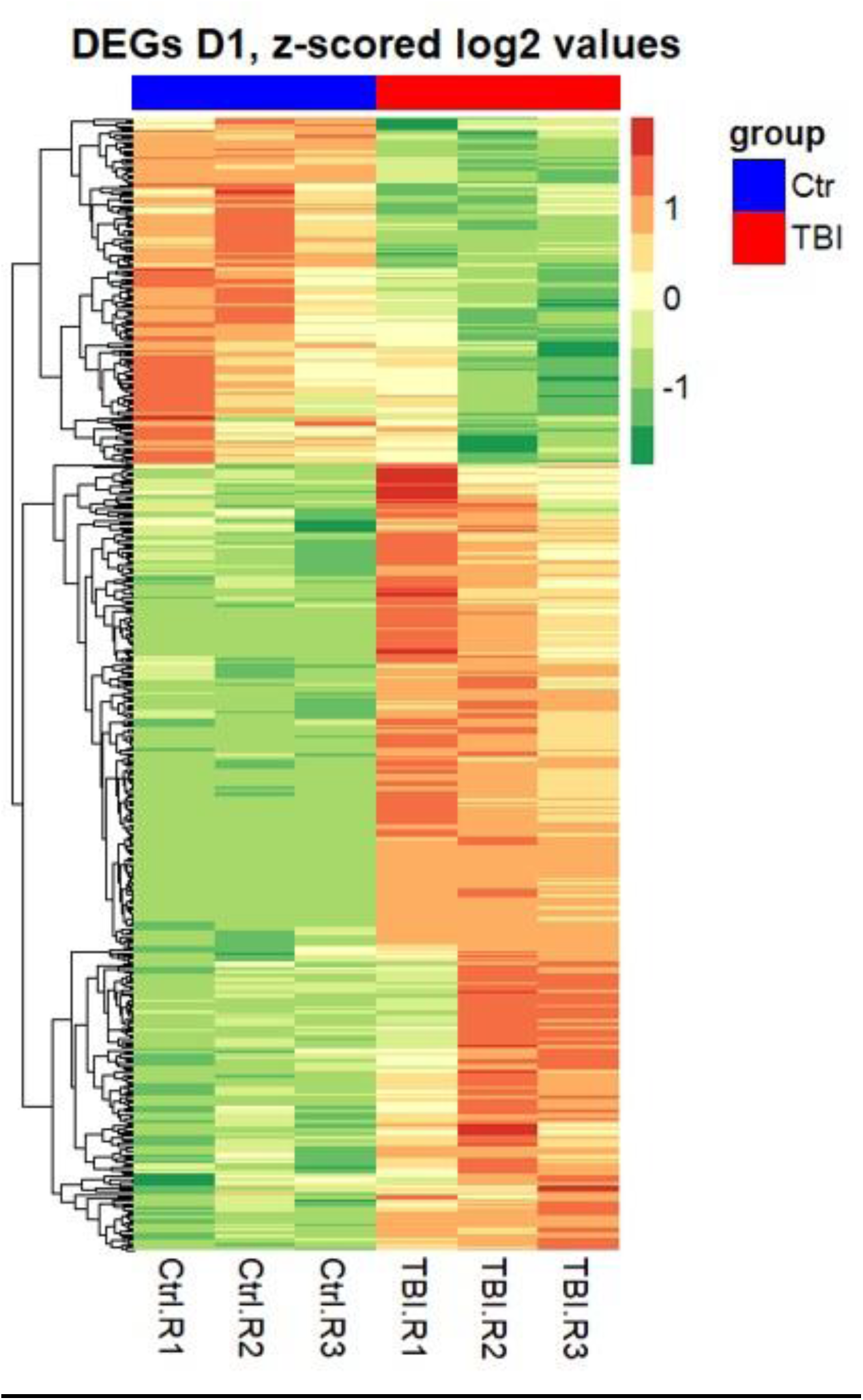
Glial gene expression heat map. Panels present clustering of differentially expressed genes (DEGs) for day 1 post-TBI. Gene expression level presented as z-scored log2(X+1) transformed values; Control replicates in blue, TBI replicates in red.

**Figure S3.**
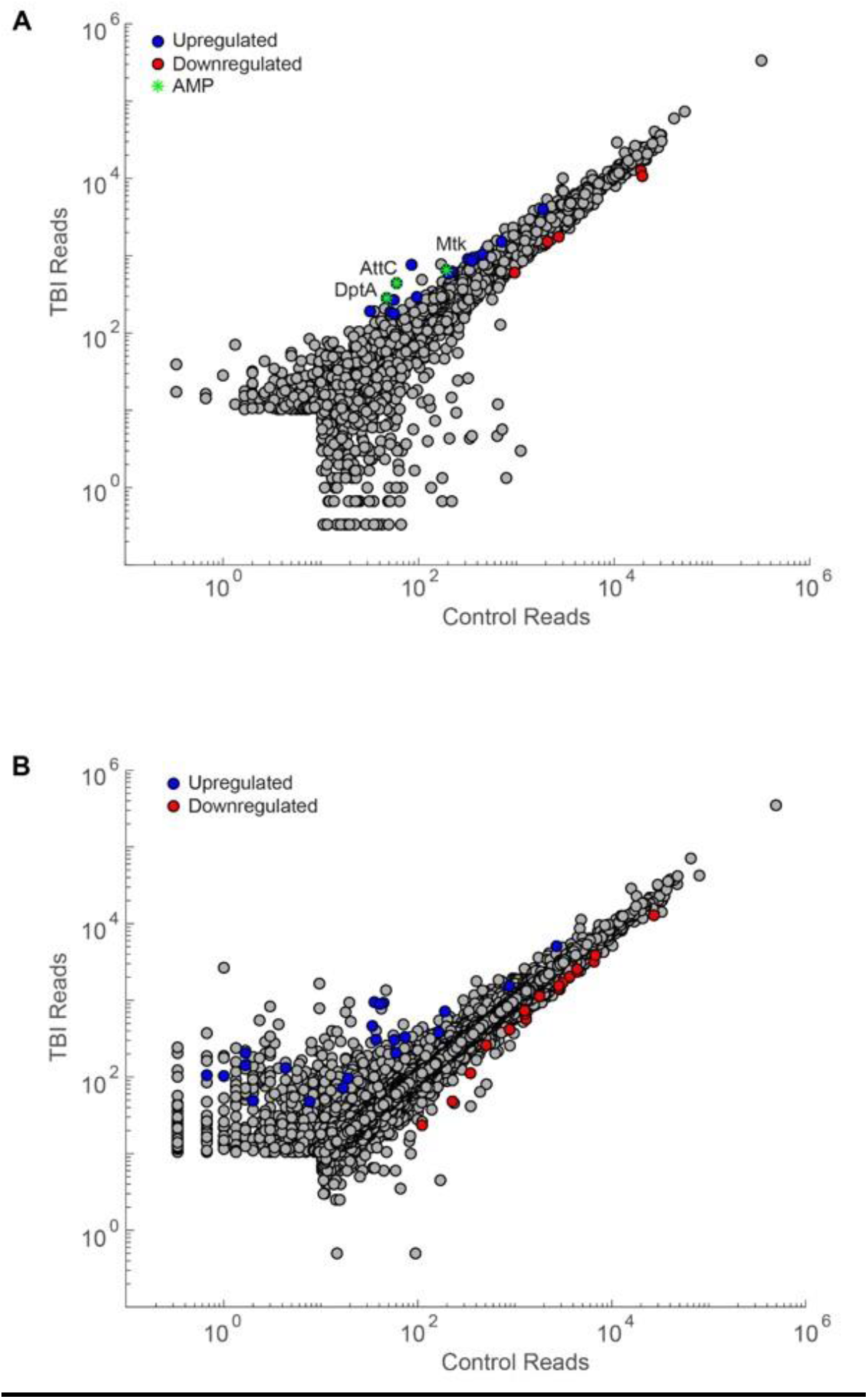
Differentially expressed genes in repo-TRAP at 3 and 7 days post-TBI. Scatter plot for glial genes where average reads in the control condition are plotted against average reads three days **(A)** or seven days **(B)** after TBI (blue dots, log_*2*_fold change ≥ 0.6, Benjamini adjusted p <0.1) or downregulated (red dots, log_*2*_fold change ≤ −0.6, Benjamini adjusted p <0.1) 24 hours after TBI induction. AMPs are indicated with green asterisks. Genes with average reads <10 in both control and TBI condition were excluded.

## Author contributions

BvA developed the TBI assay. Experiments were designed by BvA, SS and RA. Sleep, climbing and mortality assays were performed by BvA and SS. TUNEL assays were performed by SR and AR. NF-κB experiments were carried out by BvA and EB. TRAP seq was conducted by SS, FX and TQI. RNA-preprocessing, differential gene expression and functional analysis were performed by MI and RIB. BvA and RA wrote the manuscript.

## Competing interests

The authors declare no competing interests

## Code availability

The code used to generate the results that are reported in this study are available from the corresponding author upon reasonable request.

## Data availability

The data that support the findings of this study are available from the corresponding author upon reasonable request

## References

1. Maas, A.I., N. Stocchetti, and R. Bullock, Moderate and severe traumatic brain injury in adults. Lancet Neurol, 2008. 7(8): p. 728–41.

2. Menon, D.K., et al., Position statement: definition of traumatic brain injury. Arch Phys Med Rehabil, 2010. 91(11): p. 1637–40.

3. Langlois, J.A., W. Rutland-Brown, and M.M. Wald, The epidemiology and impact of traumatic brain injury: a brief overview. J Head Trauma Rehabil, 2006. 21(5): p. 375–8.

4. Sauaia, A., et al., Epidemiology of trauma deaths: a reassessment. J Trauma, 1995. 38(2): p. 185–93.

5. Gaetz, M., The neurophysiology of brain injury. Clin Neurophysiol, 2004. 115(1): p. 4–18.

6. Blennow, K., J. Hardy, and H. Zetterberg, The neuropathology and neurobiology of traumatic brain injury. Neuron, 2012. 76(5): p. 886–99.

7. Ransohoff, R.M. and B. Engelhardt, The anatomical and cellular basis of immune surveillance in the central nervous system. Nat Rev Immunol, 2012. 12(9): p. 623–35.

8. Loane, D.J. and K.R. Byrnes, Role of microglia in neurotrauma. Neurotherapeutics, 2010. 7(4): p. 366–77.

9. Morganti-Kossmann, M.C., et al., Role of cerebral inflammation after traumatic brain injury: a revisited concept. Shock, 2001. 16(3): p. 165–77.

10. Morganti-Kossmann, M.C., et al., Modulation of immune response by head injury. Injury, 2007. 38(12): p. 1392–400.

11. Davis, A.E., Mechanisms of traumatic brain injury: biomechanical, structural and cellular considerations. Crit Care Nurs Q, 2000. 23(3): p. 1–13.

12. Marklund, N., et al., Energy metabolic changes in the early post-injury period following traumatic brain injury in rats. Neurochem Res, 2006. 31(8): p. 1085–93.

13. Donat, C.K., et al., Microglial Activation in Traumatic Brain Injury. Front Aging Neurosci, 2017. 9: p. 208.

14. Russo, M.V. and D.B. McGavern, Inflammatory neuroprotection following traumatic brain injury. Science, 2016. 353(6301): p. 783–5.

15. Finnie, J.W., Neuroinflammation: beneficial and detrimental effects after traumatic brain injury. Inflammopharmacology, 2013. 21(4): p. 309–20.

16. Corps, K.N., T.L. Roth, and D.B. McGavern, Inflammation and neuroprotection in traumatic brain injury. JAMA Neurol, 2015. 72(3): p. 355–62.

17. Simon, D.W., et al., The far-reaching scope of neuroinflammation after traumatic brain injury. Nat Rev Neurol, 2017. 13(3): p. 171–191.

18. Aungst, S.L., et al., Repeated mild traumatic brain injury causes chronic neuroinflammation, changes in hippocampal synaptic plasticity, and associated cognitive deficits. J Cereb Blood Flow Metab, 2014. 34(7): p. 1223–32.

19. Morganti-Kossmann, M.C., et al., Inflammatory response in acute traumatic brain injury: a double-edged sword. Curr Opin Crit Care, 2002. 8(2): p. 101–5.

20. Jassam, Y.N., et al., Neuroimmunology of Traumatic Brain Injury: Time for a Paradigm Shift. Neuron, 2017. 95(6): p. 1246–1265.

21. Bellen, H.J., C. Tong, and H. Tsuda, 100 years of Drosophila research and its impact on vertebrate neuroscience: a history lesson for the future. Nat Rev Neurosci, 2010. 11(7): p. 514–22.

22. Marsh, J.L. and L.M. Thompson, Drosophila in the study of neurodegenerative disease. Neuron, 2006. 52(1): p. 169–78.

23. Casci, I. and U.B. Pandey, A fruitful endeavor: modeling ALS in the fruit fly. Brain Res, 2015. 1607: p. 47–74.

24. Bouleau, S. and H. Tricoire, Drosophila models of Alzheimer’s disease: advances, limits, and perspectives. J Alzheimers Dis, 2015. 45(4): p. 1015–38.

25. Lewis, E.A. and G.A. Smith, Using Drosophila models of Huntington’s disease as a translatable tool. J Neurosci Methods, 2016. 265: p. 89–98.

26. West, R.J., et al., Neurophysiology of Drosophila models of Parkinson’s disease. Parkinsons Dis, 2015. 2015: p. 381281.

27. Allen, N.J. and B.A. Barres, Neuroscience: Glia - more than just brain glue. Nature, 2009. 457(7230): p. 675–7.

28. Chung, W.S., et al., Do glia drive synaptic and cognitive impairment in disease? Nat Neurosci, 2015. 18(11): p. 1539–1545.

29. Doherty, J., et al., Ensheathing glia function as phagocytes in the adult Drosophila brain. J Neurosci, 2009. 29(15): p. 4768–81.

30. Logan, M.A. and M.R. Freeman, The scoop on the fly brain: glial engulfment functions in Drosophila. Neuron Glia Biol, 2007. 3(1): p. 63–74.

31. Logan, M.A., et al., Negative regulation of glial engulfment activity by Draper terminates glial responses to axon injury. Nat Neurosci, 2012. 15(5): p. 722–30.

32. Ziegenfuss, J.S., et al., Draper-dependent glial phagocytic activity is mediated by Src and Syk family kinase signalling. Nature, 2008. 453(7197): p. 935–9.

33. Losada-Perez, M., Glia: from ‘just glue’ to essential players in complex nervous systems: a comparative view from flies to mammals. J Neurogenet, 2018. 32(2): p. 78–91.

34. Sonnenfeld, M.J. and J.R. Jacobs, Macrophages and glia participate in the removal of apoptotic neurons from the Drosophila embryonic nervous system. J Comp Neurol, 1995. 359(4): p. 644–52.

35. Purice, M.D., et al., A novel Drosophila injury model reveals severed axons are cleared through a Draper/MMP-1 signaling cascade. Elife, 2017. 6.

36. Lemaitre, B. and J. Hoffmann, The host defense of Drosophila melanogaster. Annu Rev Immunol, 2007. 25: p. 697–743.

37. Sudmeier, L.J., et al., Persistent Activation of the Innate Immune Response in Adult Drosophila Following Radiation Exposure During Larval Development. G3 (Bethesda), 2015. 5(11): p. 2299–306.

38. Petersen, A.J., S.A. Rimkus, and D.A. Wassarman, ATM kinase inhibition in glial cells activates the innate immune response and causes neurodegeneration in Drosophila. Proc Natl Acad Sci U S A, 2012. 109(11): p. E656–64.

39. Petersen, A.J., R.J. Katzenberger, and D.A. Wassarman, The innate immune response transcription factor relish is necessary for neurodegeneration in a Drosophila model of ataxia-telangiectasia. Genetics, 2013. 194(1): p. 133–42.

40. Katzenberger, R.J., et al., A Drosophila model of closed head traumatic brain injury. Proc Natl Acad Sci U S A, 2013. 110(44): p. E4152–9.

41. Barekat, A., et al., Using Drosophila as an integrated model to study mild repetitive traumatic brain injury. Sci Rep, 2016. 6: p. 25252.

42. Huang, Y., et al., Translational profiling of clock cells reveals circadianly synchronized protein synthesis. PLoS Biol, 2013. 11(11): p. e1001703.

43. Nichols, C.D., J. Becnel, and U.B. Pandey, Methods to assay Drosophila behavior. J Vis Exp, 2012(61).

44. Shaw, P.J., et al., Correlates of sleep and waking in Drosophila melanogaster. Science, 2000. 287(5459): p. 1834–7.

45. Huber, R., et al., Sleep homeostasis in Drosophila melanogaster. Sleep, 2004. 27(4): p. 628–39.

46. Linford, N.J., et al., Measurement of lifespan in Drosophila melanogaster. J Vis Exp, 2013(71).

47. Cardillo, G. LogRank: Comparing survival curves of two groups using the log rank test. http://www.mathworks.com/matlabcentral/fileexchange/22317 2008.

48. Heiman, M., et al., Cell type-specific mRNA purification by translating ribosome affinity purification (TRAP). Nat Protoc, 2014. 9(6): p. 1282–91.

49. Nagoshi, E., et al., Dissecting differential gene expression within the circadian neuronal circuit of Drosophila. Nat Neurosci, 2010. 13(1): p. 60–8.

50. Bray, N.L., et al., Near-optimal probabilistic RNA-seq quantification. Nat Biotechnol, 2016. 34(5): p. 525–7.

51. Gramates, L.S., et al., FlyBase at 25: looking to the future. Nucleic Acids Res, 2017. 45(D1): p. D663–D671.

52. Soneson, C., M.I. Love, and M.D. Robinson, Differential analyses for RNA-seq: transcript-level estimates improve gene-level inferences. F1000Res, 2015. 4: p. 1521.

53. Love, M.I., W. Huber, and S. Anders, Moderated estimation of fold change and dispersion for RNA-seq data with DESeq2. Genome Biol, 2014. 15(12): p. 550.

54. Risso, D., et al., Normalization of RNA-seq data using factor analysis of control genes or samples. Nat Biotechnol, 2014. 32(9): p. 896–902.

55. Huang, D.W., et al., The DAVID Gene Functional Classification Tool: a novel biological module-centric algorithm to functionally analyze large gene lists. Genome Biol, 2007. 8(9): p. R183.

56. Huang, D.W., et al., DAVID Bioinformatics Resources: expanded annotation database and novel algorithms to better extract biology from large gene lists. Nucleic Acids Res, 2007. 35(Web Server issue): p. W169–75.

57. Bullard, J.H., et al., Evaluation of statistical methods for normalization and differential expression in mRNA-Seq experiments. BMC Bioinformatics, 2010. 11: p. 94.

58. Mouzon, B., et al., Repetitive mild traumatic brain injury in a mouse model produces learning and memory deficits accompanied by histological changes. J Neurotrauma, 2012. 29(18): p. 2761–73.

59. Krauss, J.K.J.J., *Movement disorders after TBI*. Brain Injury Medicine: Principles and Practice, ed. N.D.K. Zasler, D.I.; Zafonte, R.D. 2007, New York. 469–489.

60. Ustinova, K.I., et al., Physical therapy for correcting postural and coordination deficits in patients with mild-to-moderate traumatic brain injury. Physiother Theory Pract, 2015. 31(1): p. 1–7.

61. Guskiewicz, K.M., Balance assessment in the management of sport-related concussion. Clin Sports Med, 2011. 30(1): p. 89-102, ix.

62. Fujimoto, S.T., et al., Motor and cognitive function evaluation following experimental traumatic brain injury. Neurosci Biobehav Rev, 2004. 28(4): p. 365–78.

63. Hirsch, J. and L. Erlenmeyer-Kimling, Sign of taxis as a property of the genotype. Science, 1961. 134(3482): p. 835–6.

64. Feany, M.B. and W.W. Bender, A Drosophila model of Parkinson’s disease. Nature, 2000. 404(6776): p. 394–8.

65. Barone, M.C. and D. Bohmann, Assessing neurodegenerative phenotypes in Drosophila dopaminergic neurons by climbing assays and whole brain immunostaining. J Vis Exp, 2013(74): p. e50339.

66. Ali, Y.O., et al., Assaying locomotor, learning, and memory deficits in Drosophila models of neurodegeneration. J Vis Exp, 2011(49).

67. McCall, K., J.S. Peterson, and T.L. Pritchett, Detection of cell death in Drosophila. Methods Mol Biol, 2009. 559: p. 343–56.

68. Sun, Y., et al., TRPA channels distinguish gravity sensing from hearing in Johnston’s organ. Proc Natl Acad Sci U S A, 2009. 106(32): p. 13606–11.

69. Kamikouchi, A., et al., The neural basis of Drosophila gravity-sensing and hearing. Nature, 2009. 458(7235): p. 165–71.

70. Mettang, M., et al., IKK2/NF-kappaB signaling protects neurons after traumatic brain injury. FASEB J, 2018. 32(4): p. 1916–1932.

71. Sandsmark, D.K., J.E. Elliott, and M.M. Lim, Sleep-Wake Disturbances After Traumatic Brain Injury: Synthesis of Human and Animal Studies. Sleep, 2017. 40(5).

72. Willie, J.T., et al., Controlled cortical impact traumatic brain injury acutely disrupts wakefulness and extracellular orexin dynamics as determined by intracerebral microdialysis in mice. J Neurotrauma, 2012. 29(10): p. 1908–21.

73. Lim, M.M., et al., Dietary therapy mitigates persistent wake deficits caused by mild traumatic brain injury. Sci Transl Med, 2013. 5(215): p. 215ra173.

74. Rowe, R.K., et al., Diffuse brain injury induces acute post-traumatic sleep. PLoS One, 2014. 9(1): p. e82507.

75. Morawska, M.M., et al., Sleep Modulation Alleviates Axonal Damage and Cognitive Decline after Rodent Traumatic Brain Injury. J Neurosci, 2016. 36(12): p. 3422–9.

76. Thomasy, H., Tumor Necrosis Factor alpha as a Potential Mediator of the Effects of Phosphodiesterase 4B Inhibition on Cognition after Traumatic Brain Injury. J Neurosci, 2016. 36(46): p. 11587–11589.

77. Hazra, A., et al., Delayed thalamic astrocytosis and disrupted sleep-wake patterns in a preclinical model of traumatic brain injury. J Neurosci Res, 2014. 92(11): p. 1434–45.

78. Petraglia, A.L., et al., The spectrum of neurobehavioral sequelae after repetitive mild traumatic brain injury: a novel mouse model of chronic traumatic encephalopathy. J Neurotrauma, 2014. 31(13): p. 1211–24.

79. Skopin, M.D., et al., Chronic decrease in wakefulness and disruption of sleep-wake behavior after experimental traumatic brain injury. J Neurotrauma, 2015. 32(5): p. 289– 96.

80. Thomas, J., et al., Identification of an intronic splicing regulatory element involved in auto-regulation of alternative splicing of SCL33 pre-mRNA. Plant J, 2012. 72(6): p. 935– 46.

81. Xiong, W.C., et al., repo encodes a glial-specific homeo domain protein required in the Drosophila nervous system. Genes Dev, 1994. 8(8): p. 981–94.

82. Huang, Y., F.S. Ng, and F.R. Jackson, Comparison of larval and adult Drosophila astrocytes reveals stage-specific gene expression profiles. G3 (Bethesda), 2015. 5(4): p. 551–8.

83. Auld, V.J., et al., Gliotactin, a novel transmembrane protein on peripheral glia, is required to form the blood-nerve barrier in Drosophila. Cell, 1995. 81(5): p. 757–67.

84. Bainton, R.J., et al., moody encodes two GPCRs that regulate cocaine behaviors and blood-brain barrier permeability in Drosophila. Cell, 2005. 123(1): p. 145–56.

85. Benjamini, Y.H., Y, Controlling the false discovery rate: a practical and powerful approach to multiple testing. Journal of the Royal Statistical Society, Series B, 1995. 57(1): p. 289–300.

86. Ekengren, S. and D. Hultmark, A family of Turandot-related genes in the humoral stress response of Drosophila. Biochem Biophys Res Commun, 2001. 284(4): p. 998–1003.

87. Imler, J.L. and P. Bulet, Antimicrobial peptides in Drosophila: structures, activities and gene regulation. Chem Immunol Allergy, 2005. 86: p. 1–21.

88. Ng, F.S., et al., TRAP-seq Profiling and RNAi-Based Genetic Screens Identify Conserved Glial Genes Required for Adult Drosophila Behavior. Front Mol Neurosci, 2016. 9: p. 146.

89. Buchon, N., N. Silverman, and S. Cherry, Immunity in Drosophila melanogaster-from microbial recognition to whole-organism physiology. Nat Rev Immunol, 2014. 14(12): p. 796–810.

90. De Gregorio, E., et al., Genome-wide analysis of the Drosophila immune response by using oligonucleotide microarrays. Proc Natl Acad Sci U S A, 2001. 98(22): p. 12590–5.

91. Cerenius, L. and K. Soderhall, Coagulation in invertebrates. J Innate Immun, 2011. 3(1): p. 3–8.

92. Kambris, Z., et al., Drosophila immunity: a large-scale in vivo RNAi screen identifies five serine proteases required for Toll activation. Curr Biol, 2006. 16(8): p. 808–13.

93. Lemaitre, B., et al., The dorsoventral regulatory gene cassette spatzle/Toll/cactus controls the potent antifungal response in Drosophila adults. Cell, 1996. 86(6): p. 973– 83.

94. Verma, R.P. and C. Hansch, Matrix metalloproteinases (MMPs): chemical-biological functions and (Q)SARs. Bioorg Med Chem, 2007. 15(6): p. 2223–68.

95. Welch, W.J., Mammalian stress response: cell physiology, structure/function of stress proteins, and implications for medicine and disease. Physiol Rev, 1992. 72(4): p. 1063–81.

96. Chen, J., et al., Participation of the p38 pathway in Drosophila host defense against pathogenic bacteria and fungi. Proc Natl Acad Sci U S A, 2010. 107(48): p. 20774–9.

97. Kume, K., et al., Dopamine is a regulator of arousal in the fruit fly. J Neurosci, 2005. 25(32): p. 7377–84.

98. Wu, M.N., et al., A genetic screen for sleep and circadian mutants reveals mechanisms underlying regulation of sleep in Drosophila. Sleep, 2008. 31(4): p. 465–72.

99. Ueno, T. and K. Kume, Functional characterization of dopamine transporter in vivo using Drosophila melanogaster behavioral assays. Front Behav Neurosci, 2014. 8: p. 303.

100. Pearson, J.C., et al., Multiple transcription factor codes activate epidermal wound-response genes in Drosophila. Proc Natl Acad Sci U S A, 2009. 106(7): p. 2224–9.

101. Gaudet, P.L.M.; Thomas, P., Gene Ontology annotation inferences using phylogenetic trees. GO Reference Genome Project, 2010. http://www.geneontology.org/cgi-bin/references.cgi#GO_REF0000033.

102. Oeckinghaus, A., M.S. Hayden, and S. Ghosh, Crosstalk in NF-kappaB signaling pathways. Nat Immunol, 2011. 12(8): p. 695–708.

103. Mattson, M.P., NF-kappaB in the survival and plasticity of neurons. Neurochem Res, 2005. 30(6-7): p. 883–93.

104. Helmy, A., et al., The cytokine response to human traumatic brain injury: temporal profiles and evidence for cerebral parenchymal production. J Cereb Blood Flow Metab, 2011. 31(2): p. 658–70.

105. Bales, K.R., et al., The NF-kappaB/Rel family of proteins mediates Abeta-induced neurotoxicity and glial activation. Brain Res Mol Brain Res, 1998. 57(1): p. 63–72.

106. Brambilla, R., et al., Inhibition of astroglial nuclear factor kappaB reduces inflammation and improves functional recovery after spinal cord injury. J Exp Med, 2005. 202(1): p. 145–56.

107. Yang, K., X.S. Mu, and R.L. Hayes, Increased cortical nuclear factor-kappa B (NF-kappa DNA binding activity after traumatic brain injury in rats. Neurosci Lett, 1995. 197(2): p. 101–4.

108. Nonaka, M., et al., Prolonged activation of NF-kappaB following traumatic brain injury in rats. J Neurotrauma, 1999. 16(11): p. 1023–34.

109. Hu, Y.C., et al., Biphasic activation of nuclear factor kappa B and expression of p65 and c-Rel after traumatic brain injury in rats. Inflamm Res, 2014. 63(2): p. 109–15.

110. Khush, R.S., F. Leulier, and B. Lemaitre, Drosophila immunity: two paths to NF-kappaB. Trends Immunol, 2001. 22(5): p. 260–4.

111. Kuo, T.H., et al., Sleep triggered by an immune response in Drosophila is regulated by the circadian clock and requires the NFkappaB Relish. BMC Neurosci, 2010. 11: p. 17.

112. Cao, Y., et al., Dnr1 mutations cause neurodegeneration in Drosophila by activating the innate immune response in the brain. Proc Natl Acad Sci U S A, 2013. 110(19): p. E1752–60.

113. Kounatidis, I., et al., NF-kappaB Immunity in the Brain Determines Fly Lifespan in Healthy Aging and Age-Related Neurodegeneration. Cell Rep, 2017. 19(4): p. 836–848.

114. Hayden, M.S. and S. Ghosh, NF-kappaB, the first quarter-century: remarkable progress and outstanding questions. Genes Dev, 2012. 26(3): p. 203–34.

115. Hetru, C. and J.A. Hoffmann, NF-kappaB in the immune response of Drosophila. Cold Spring Harb Perspect Biol, 2009. 1(6): p. a000232.

116. Sun, M. and L.L. Chen, A Novel Method to Model Chronic Traumatic Encephalopathy in Drosophila. J Vis Exp, 2017(125).

117. Bartholomew, N.R., et al., Impaired climbing and flight behaviour in Drosophila melanogaster following carbon dioxide anaesthesia. Sci Rep, 2015. 5: p. 15298.

118. Statler, K.D., et al., Isoflurane improves long-term neurologic outcome versus fentanyl after traumatic brain injury in rats. J Neurotrauma, 2000. 17(12): p. 1179–89.

119. Statler, K.D., et al., Comparison of seven anesthetic agents on outcome after experimental traumatic brain injury in adult, male rats. J Neurotrauma, 2006. 23(1): p. 97–108.

120. Statler, K.D., et al., Isoflurane exerts neuroprotective actions at or near the time of severe traumatic brain injury. Brain Res, 2006. 1076(1): p. 216–24.

121. Fischer, J.A., et al., Anesthetics Influence Mortality in a Drosophila Model of Blunt Trauma With Traumatic Brain Injury. Anesth Analg, 2018. 126(6): p. 1979–1986.

122. Graber, D.J., B.A. Costine, and W.F. Hickey, Early inflammatory mediator gene expression in two models of traumatic brain injury: ex vivo cortical slice in mice and in vivo cortical impact in piglets. J Neuroinflammation, 2015. 12: p. 76.

123. Morganti, J.M., L.K. Riparip, and S. Rosi, Call Off the Dog(ma): M1/M2 Polarization Is Concurrent following Traumatic Brain Injury. PLoS One, 2016. 11(1): p. e0148001.

124. Lagraoui, M., et al., Controlled cortical impact and craniotomy induce strikingly similar profiles of inflammatory gene expression, but with distinct kinetics. Front Neurol, 2012. 3: p. 155.

125. Meng, Q., et al., Traumatic Brain Injury Induces Genome-Wide Transcriptomic, Methylomic, and Network Perturbations in Brain and Blood Predicting Neurological Disorders. EBioMedicine, 2017. 16: p. 184–194.

126. Almeida-Suhett, C.P., et al., Temporal course of changes in gene expression suggests a cytokine-related mechanism for long-term hippocampal alteration after controlled cortical impact. J Neurotrauma, 2014. 31(7): p. 683–90.

127. Williams, J.A., et al., Interaction between sleep and the immune response in Drosophila: a role for the NFkappaB relish. Sleep, 2007. 30(4): p. 389–400.

128. Kuo, T.H., A. Handa, and J.A. Williams, Quantitative measurement of the immune response and sleep in Drosophila. J Vis Exp, 2012(70): p. e4355.

129. Nappi, A.J. and E. Vass, Melanogenesis and the generation of cytotoxic molecules during insect cellular immune reactions. Pigment Cell Res, 1993. 6(3): p. 117–26.

